# A chimeric viral platform for directed evolution in mammalian cells

**DOI:** 10.1101/2024.04.20.590384

**Authors:** Alexander J. Cole, Christopher E. Denes, Cesar L. Moreno, Thomas Dobson, Daniel Hesselson, G. Gregory Neely

## Abstract

Directed evolution (DE) is a process of mutation and artificial selection to breed biomolecules with new or improved activity^1,2^. DE platforms are primarily prokaryotic or yeast-based, and stable mutagenic mammalian systems have been challenging to establish and apply^3^. To this end, we developed PROTein Evolution Using Selection (PROTEUS), a new platform that uses chimeric virus-like vesicles (VLVs) to enable extended mammalian DE campaigns without loss of system integrity. This platform is stable and can generate sufficient diversity for DE in mammalian systems. Using PROTEUS, we altered the doxycycline responsiveness of tetracycline-controlled transactivators, generating a more sensitive TetON-4G tool for gene regulation. PROTEUS is also compatible with intracellular nanobody evolution, and we use it to design a DNA damage-responsive anti-p53 nanobody. Overall, PROTEUS is an efficient and stable platform to direct evolution of biomolecules within mammalian cells.

## Main

Using iterative rounds of diversification, selection, and amplification, directed evolution (DE) can produce biomolecules with new or improved functions^1,2,4–7^. While this approach has been widely used to evolve molecules in simple prokaryotic and eukaryotic systems^8,9^, these environments lack the full complement of post-translational modifications, protein-protein interactions, and signaling networks found in mammalian cells^3^.

Ideally, proteins destined for mammalian applications would be evolved directly in mammalian cells. Historically, this has been achieved using ex mammalia mutagenesis techniques combined with phenotypic screening in mammalian cells^10^. More recently, targeted mutagenesis has linked protein function to selectable or screenable markers allowing target diversification and variant selection in the same mammalian cell^11,12^. However, cell-based approaches that link an integrated target molecule to cellular fitness can be derailed by mutations in the host genome^13^. Placing the target in a viral genome can mitigate this issue since naive host cells can be provided for each round of DE. However, existing virus-based mammalian DE systems are limited by safety concerns^14^, low mutational rates^15^, are target-specific^16,17^, or lack functionality^18,19^.

Here, we describe PROTein Evolution Using Selection (PROTEUS), a new platform that uses chimeric virus-like vesicles (VLVs) to enable extended mammalian DE campaigns without loss of system integrity. PROTEUS rapidly generates authentic evolution products with superior functionality in mammalian cells, and will have broad utility for evolving proteins *in situ*.

### Capsid-deficient VLVs support host-dependent propagation of the SFV genome

Alphavirus genomic RNA (gRNA) is recognized by a strain-specific capsid protein that packages gRNA into infectious particles. While cognate packaging signals encoded in the gRNA are sufficient for encapsidation^20^, there are additional redundant packaging sequences distributed throughout the alphavirus genome^21^. In the context of DE, these interactions generate “cheater” particles that interfere with viral replication and contribute to a failure to recover authentic DE products^19^. While an intact capsid is essential for the pathogenicity of blood-borne viruses, the capsid protein is dispensable for *in vitro* propagation of VLVs^22^.

To explore whether eliminating capsid-gRNA interactions enables robust host-dependent viral propagation, we designed a chimeric two-component system based on a self-replicating Semliki Forest Virus (SFV) replicon^23^, where the infectivity of SFV VLVs is determined by the expression level of the Indiana vesiculovirus G (VSVG) coat protein in a host mammalian cell permissive to SFV replication (BHK-21, **Fig. 1a**). Importantly, VSVG does not encapsidate viral RNA and there are no regions of significant homology between the RNA encoding VSVG and the SFV genome, reducing opportunities for recombination events that could restore replication competence. We generated an SFV DNA replicon incorporating fourteen point mutations (four synonymous) in the Non-Structural Proteins (NSPs 1-4), which were reported to produce high-titer SFV VLVs pseudotyped with VSVG (**Extended Data Fig. 1a**,^24^). Further, to reduce the cytopathic effects of SFV transduction, we exchanged a three amino acid loop within NSP2 with an attenuated variant (A674R/D675L/A676E^25^) to generate the pSFV-DE replicon construct (**Fig. 1A, Extended Data Fig. 1a**). Attenuation did not affect SFV-DE/VSVG VLV titer or amplification factor (the ratio of VLVs released per VLV transduced) (**Extended Data Fig. 1b and c**), indicating that reduced cytotoxicity was achieved without compromising VLV fitness.

**Fig. 1.**
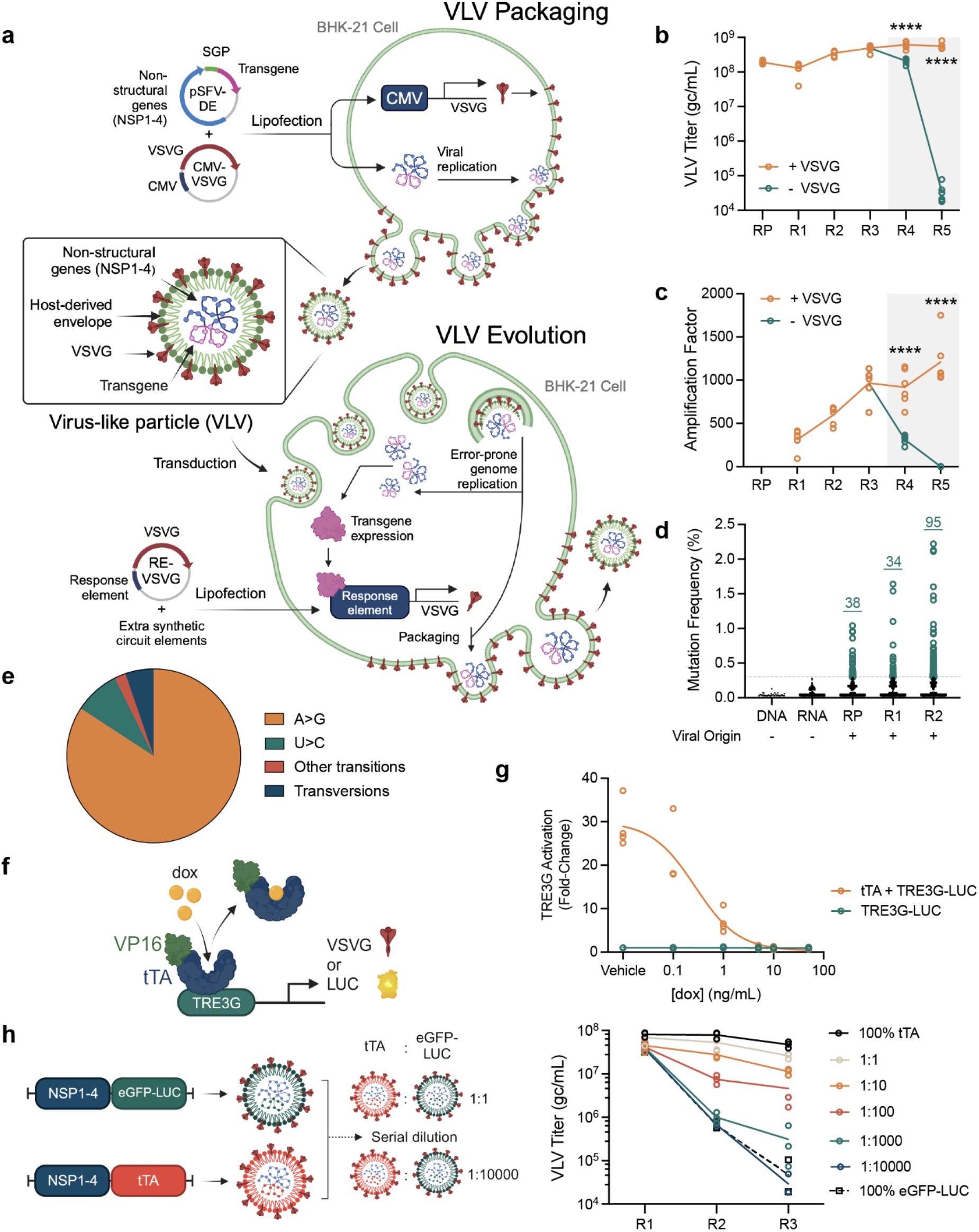
Host-dependent propagation of VLVs. (**a**) SFV VLVs are initially packaged using SFV-DE DNA replicons that encode the DE target transgene in cells that constitutively express VSVG. Infectious VLVs are propagated for evolution in host cells that express VSVG under the control of a circuit that is directly or indirectly regulated by the DE target. Titers (**b**) and amplification factors (**c**) of eGFP-LUC VLVs propagated in cells constitutively expressing CMV_VSVG (+VSVG) for all rounds (R) RP-R5 or control DNA (-VSVG; for R4 and R5 presented within the gray box) (*N* = 6). (**d**) Allele frequency of mutations in a neutral eGFP transgene. Dotted line represents detection limit for viral variants (0.3%). (**e**) Mutational spectrum of viral variants from R2 (*N* = 95). (**f**) Circuit design for tTA mediated activation of TRE3G. (**g**) Dox-dependent repression of a TRE3G-regulated LUC reporter (*N* = 4). (**h**) Titers of serially diluted tTA VLVs propagated on cells expressing VSVG under the control of TRE3G in the absence of doxycycline (*N* = 3). Colors are for illustrative purposes and do not reflect different membrane compositions.

Implementation of chimeric VLVs for mammalian DE campaigns requires complementation by host cell expression of VSVG. To confirm host-dependence, SFV-DE VLVs carrying an eGFP-P2A-Luciferase (eGFP-LUC) reporter were propagated for multiple rounds at high titer (>10^8^ genome copies (gc)/mL; **Fig. 1b**, R1-R3) and then used to infect VSVG- or mock-transfected host cells (**Fig. 1b**, R4-R5). VSVG-transfected cells supported SFV-DE VLV titers leading to an amplification factor >1000 (**Fig. 1c**). Without VSVG, however, VLV titers drop below the limit of detection and these mock-transfected (VSVG negative) cells showed an amplification factor <1.

While eGFP expression was maintained in cells that constitutively expressed VSVG (**Extended Data Fig. 1d**), no eGFP expression was detectable after two rounds of transduction on mock-transfected cells (**Extended Data Fig. 1e**). Of note, VSVG-expressing cells showed a gradual reduction in eGFP levels over 5 rounds of transduction with SFV-DE VLVs (**Extended Data Fig. 1d**). Since VSVG levels were constitutive and not dependent on the viral transgene, and since smaller viral genomes have a replicative advantage^26^, we hypothesized that the reduced eGFP signal was due to truncation of the GFP transgene over rounds of transduction. Indeed, rounds of transduction were accompanied by progressive truncation of the viral transgene (**Extended Data Fig. 1f**), suggesting that selective pressure is required to maintain a full-length viral transgene.

DE requires diversification of the target transgene to produce variants with increased fitness. Alphaviruses are error-prone with reported mutation frequencies >10^−4^ per nucleotide in each round of replication^27^. Using non-viral DNA and RNA templates, we established a detection limit of 0.3% for new mutations by amplicon deep sequencing (dotted line, **Fig. 1d**). We observed an accumulation of bona fide mutations (∼1 per 10^4^ transduced cells) during the propagation of VLVs carrying the unselected eGFP-LUC reporter (**Fig. 1d**, RP-R2). Multiple substitution types were observed with a strong A-to-G (and complementary U-to-C) transition mutational bias (**Fig. 1e, Supplementary Table 1**, ^28^).

Application of selective pressure to a target transgene requires a tight link between the resulting expression level of VSVG and viral infectivity (i.e. fitness). To test whether transgene-dependent VSVG induction gives VLVs a selective advantage, we used a circuit that is activated by the tetracycline-controlled transactivator (tTA; **Fig. 1f**) in the absence of doxycycline (dox) (**Fig. 1g**). VLVs carrying the circuit-activating tTA transgene were serially diluted with VLVs carrying a neutral eGFP-LUC transgene and propagated in the absence of dox (**Fig. 1h**). Within 3 rounds, the titers of populations containing tTA VLVs diverged from the non-activating eGFP-LUC control (**Fig. 1h**). This was consistent with the propagation-associated enrichment of the tTA transgene at dilutions up to 1:1000 (**Extended Data Fig. 2**). We next tested a serum-responsive circuit (**Extended Data Fig. 3a and b**,^19^) in which expression of the serum response factor DNA binding domain (SRF[DBD]) fused to the VP64 activation domain (SRF-VP64) further enhanced circuit activity (**Extended Data Fig. 3c**). VLVs carrying the circuit-activating SRF-VP64 transgene exhibited a small proliferative advantage over those carrying eGFP-LUC (**Extended Data Fig. 3d and e**), which correlated with increasing prevalence in a direct competition experiment within four rounds (**Extended Data Fig. 3f-h**). Selection rapidly favored a shorter SRF transgene that retained a minimal DNA-binding domain^29^ (**Extended Data Fig. 3i**). Chimeric VLVs do not indiscriminately package significant amounts of VSVG RNA, and none was detected by R4 (**Extended Data Fig. 3j**), indicating that VLV propagation is completely dependent on host cell expression of VSVG. PROTEUS, therefore, resolves problems identified in other virus-based mammalian DE systems^19^.

### PROTEUS generates authentic evolution products

To test whether a circuit linking a VLV-encoded transgene to VSVG production can evolve a new protein function, we used a tTA-regulated circuit (**Fig. 1f**) to select for doxycycline (dox) resistance in tTA. Saturating clonal selections in bacteria have identified point mutations distributed across the tTA protein that confer dox resistance^30^, some of which are also sufficient to provide dox resistance in mammalian cells^15^. Having confirmed that tTA activity was suppressed by dox in BHK-21 cells using a luciferase reporter under the control of an optimized tetracycline response element (TRE3G^31^; **Fig. 1g**), we asked whether tTA activity could support VLV propagation on a TRE3G-regulated VSVG circuit under mild selective pressure. tTA and eGFP-LUC VLVs were independently packaged and amplified in cells constitutively expressing VSVG before switching to a TRE3G-regulated circuit exposed to minimally inhibiting dox (0.1 ng/mL, E1-E3, gray box), where tTA provided a large selective advantage over the neutral eGFP-LUC transgene (**Fig. 2a and b**). Note, all VLVs amplified in cells constitutively expressing VSVG show efficient transduction at E1 so the exponential selective advantage of circuit activation is observed from round E2.

**Fig. 2.**
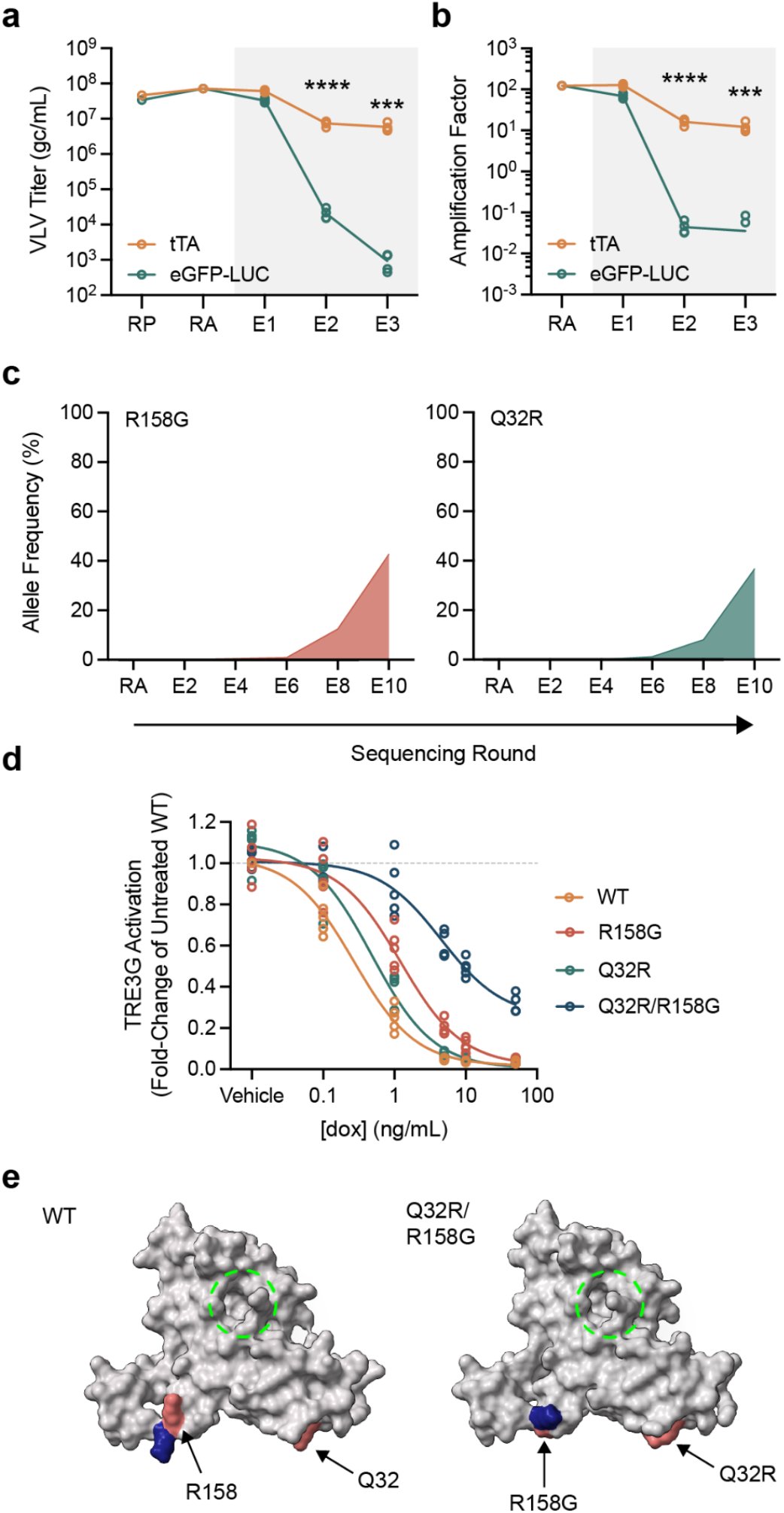
PROTEUS generates authentic evolution products. (**a**) Titers and amplification factors of VLVs propagated on cells expressing VSVG under the control of TRE3G (gray box) (*N* = 4). RP (packaging) and RA (amplification) indicate rounds of VLV propagation under constitutive VSVG expression, while E1-X labelling indicates rounds of evolution under transgene-regulated VSVG expression. (**b**) Allele frequency of the major variants identified in Campaign 1. (**c**) Dox-resistance of evolved tTA variants (*N* = 5). (**d**) Variant-induced structural changes in tTA modelled with AlphaFold2 (red, mutated residues; blue, displaced functional groups; dashed green circle, drug binding pocket).

We conducted two independent ten-round DE campaigns to isolate dox-resistant tTA variants (**Extended Data Fig. 4a**). A R158G variant (resulting from an A-to-G transition (**Fig. 1e**)) was detected in both campaigns by E4 (**Supplementary Table 2**), with different second-site mutations appearing by E6: Q32R (Campaign 1, **Fig. 2c**) and a triple mutant D178G/H179R/Q180R (Campaign 2, **Extended Data Fig. 4b and c**). Both variants from Campaign 1 have been previously identified as dox resistance mutants^30,32^. In isolation, R158G provided modest dox resistance, while Q32R had minimal effect (**Fig. 2d, Supplementary Table 3**). However, the Q32R/R158G double mutant showed strong resistance to fully inhibitory concentrations of dox (**Fig. 2d**), consistent with their synchronized increase in frequency once both appeared in the population (**Fig. 2c, Supplementary Table 2**). Similarly, in Campaign 2, the DHQ-to-GRR mutation enhanced the dox resistance of the initial R158G mutation (**Extended Data Fig. 4d, Supplementary Table 3**). Within this cluster of mutations at positions 178-180, all of the dox resistance was associated with the previously identified D178G variant (**Extended Data Fig. 4e**,^30^), indicating that H179R/Q180R were passengers arising from a complex mutational event. Structural modelling of the parental tTA protein with AlphaFold2^33^ was congruent with the drug-bound crystal structure (**Extended Data Fig. 4f**). Therefore, we modelled the dox resistant mutations from both campaigns, revealing side chain rearrangements in a linker region distal to the drug-binding site (**Fig. 2e, Extended Data Fig. 4g**). Together, these campaigns recovered authentic evolution products in tTA, validating SFV-DE VLVs as an efficient platform for PROTein Evolution Using Selection (PROTEUS).

### Enhancing drug-inducible transcriptional control

We next asked if PROTEUS can be used to improve existing molecular tools. Here, we focused on increasing the dox sensitivity of the third-generation reverse tetracycline-controlled transactivator (rtTA-3G), which has been extensively optimized using other methods^14,34^. Using a TRE3G-regulated circuit (**Fig. 3a**), the parental rtTA-3G protein had an EC50 of 39 ng/mL dox (**Fig. 3b**). On the TRE3G-regulated circuit, rtTA-3G VLVs had a large fitness advantage over neutral eGFP-LUC VLVs at 100 ng/mL dox (**Fig. 3c, Extended Data Fig. 5a**). We propagated rtTA-3G VLVs for 30 rounds, adjusting the dox concentration to maintain strong selective pressure for increased dox sensitivity (**Extended Data Fig. 5b**). The VLVs were sequenced every 5 rounds (**Extended Data Fig. 5c**), revealing an M59I variant that appeared early in the campaign and reached fixation by E30 (**Fig 3d, Supplementary Table 4**). A second D5N variant appeared by E20 and the majority of rtTA-3G transgenes (57.75%) carried both mutations by E30 (**Fig 3d, Extended Data Fig. 5d, Supplementary Table 4**). No additional variants were detected by E60 at 3 ng/mL dox suggesting that this campaign reached a local fitness peak. Both single mutations individually enhanced the dox sensitivity of rtTA-3G and the D5N/M59I double mutant was further improved (EC50 7 ng/mL; **Fig. 3e, Supplementary Table 3**), without increased leakiness in the absence of dox (**Extended Data Fig. 5e**). Both mutations were distal to the dox binding site and were not predicted to cause any major structural changes (**Extended Data Fig. 5f**). Overall, the D5N/M59I variant has a superior dox response profile and will find broad utility as a fourth generation rtTA tool (rtTA-4G) in challenging applications that require tight control of gene expression.

**Fig. 3.**
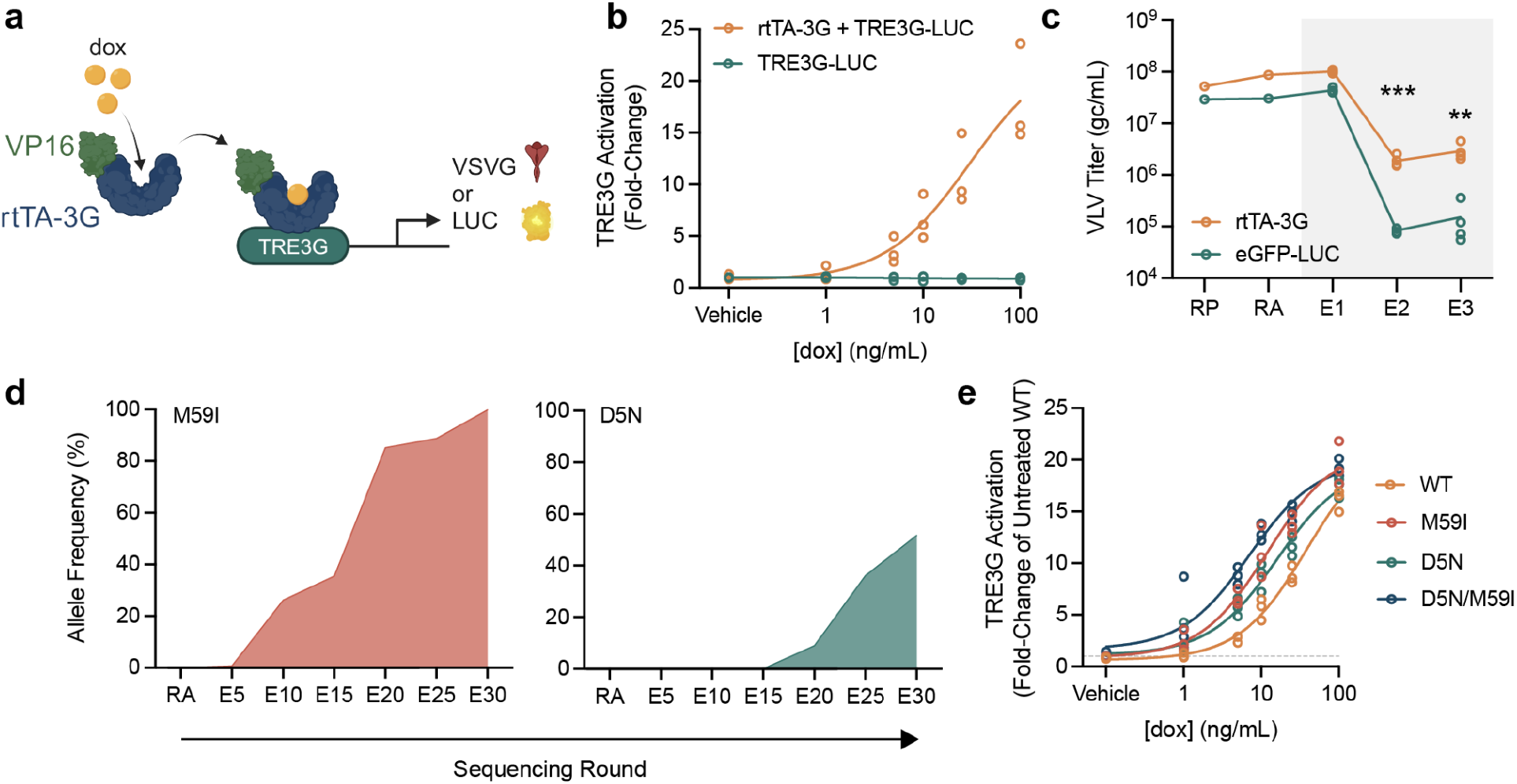
Enhancing drug-inducible transcriptional control. (**a**) Circuit design for rtTA-mediated activation of TRE3G. (**b**) Dox-dependent activation of a TRE3G-regulated LUC reporter (*N* = 3). (**c**) Titers of VLVs propagated on cells expressing VSVG under the control of TRE3G at 100 ng/mL dox (gray box) (*N* = 4). (**d**) Allele frequency of the major variants identified during long-term propagation on minimal concentrations of dox. (**e**) Dox-sensitivity of evolved tTA variants (*N* = 3).

### Directed evolution of an intracellular nanobody

Intracellular nanobodies (Nb) have potential for interrogating or modulating “undruggable” targets but are prone to instability when localized inside mammalian cells^35^. For example, a p53 biosensor based on Nb139 (**Extended Data Fig. 6a**,^36^) does not localize to the nucleus in response to cisplatin (**Extended Data Fig. 6b**) despite robust nuclear p53 accumulation (**Extended Data Fig. 6c**). To improve Nb139 interaction with p53, we conducted a DE campaign regulated by Nb139 activating a p53-dependent synthetic circuit. Using a 2-hybrid circuit design with a p53 bait (**Fig. 4a**), the parental Nb139-VP64 fusion successfully activated a reporter circuit (**Fig. 4b**), and Nb139-VP64 VLVs outcompeted neutral eGFP-LUC VLVs (**Fig 4c, Extended Data Fig. 6d**). Long-term propagation of Nb139-VP64 VLVs (**Extended Data Fig. 6e**) led to the accumulation of S26P and Y60C mutations that evolved to fixation in the population (**Fig. 4d, Extended Data Fig. 6f, Supplementary Table 5**). S26P and Y60C map to framework region (FR) 1 and FR2, respectively, indicating that these variant residues do not interact directly with p53 (**Fig. 4e**). However, only the S26P variant increased reporter activity at early and late timepoints (**Fig. 4f**), with no further increase observed in the S26P/Y60C double mutant. To test whether S26P improved the sensitivity of a p53 biosensor (**Extended Data Fig. 6a**), we expressed Nb139-eGFP fusions (parental and variants) in BHK-21 cells. In response to cisplatin, Nb139[S26P]-eGFP translocated to the nucleus and labeled nuclear puncta prior to cell death, in contrast to uniform expression throughout the cell in tGFP controls (**Fig. 4g, Extended Data Fig. 7, Supplementary Movie 1**). The S26P single mutant exhibited the largest response to cisplatin, while Y60C also modestly improved sensitivity in the biosensor format (**Fig. 4h**). This campaign demonstrates that the intracellular function of Nb139 (crystal structure 4QO1), which was already classified as stable^35^, can be further improved through evolution within a mammalian cell. Notably, the evolved variants did not directly alter the Nb139-p53 binding interface. Overall, our application of PROTEUS here has generated a novel p53 biosensor that allows the visualization of p53 nuclear recruitment *in vivo*.

**Fig. 4.**
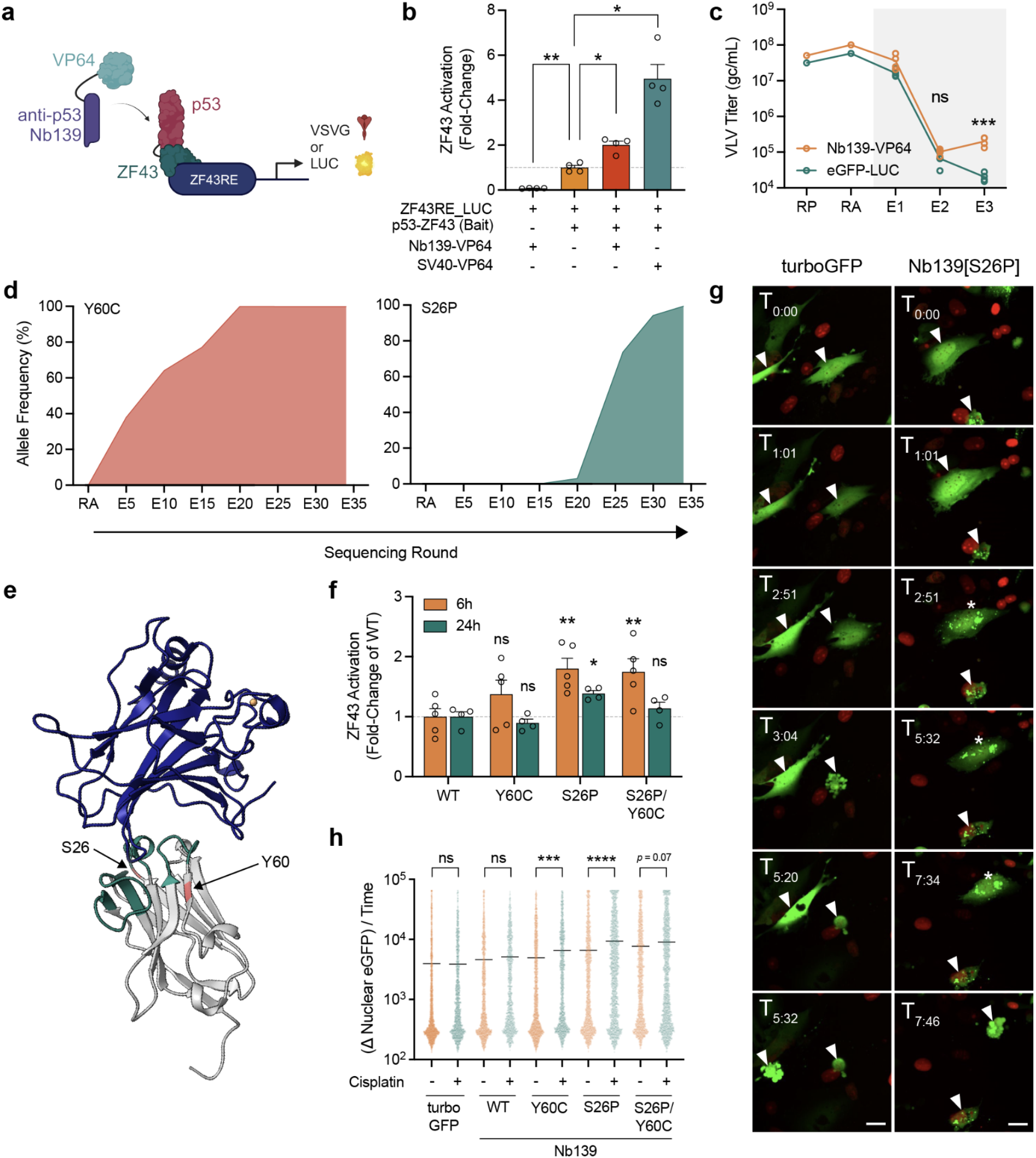
Directed evolution of an intracellular nanobody. (**a**) 2-hybrid circuit design for nanobody-p53 interactions. (**b**) Recruitment of Nb139-VP64 to a p53 bait activates the 2-hybrid circuit (SV40, positive interaction control (PMID: 9043710, *N* = 4)). Normalized to the ZF43RE_LUC + p53-ZF43 bait incomplete circuit. (**c**) Titers of VLVs propagated on cells expressing VSVG under the control of a p53 2-hybrid circuit (gray box) (*N* = 4). (**d**) Allele frequency of the major variants identified during long-term propagation on the 2-hybrid circuit. (**e**) Crystal structure 4QO1 showing Nb139 (gray; green, Nb139 complementarity-determining regions; red, evolved variant positions) in a complex with p53 (blue). (**f**) Circuit activation by Nb139-VP64 variants in cells expressing a p53 bait and LUC reporter (*N* = 4). (**g**) Timelapse of cisplatin-treated cells that express Nb139[S26P]-eGFP fusion or GFP alone (red, nuclei labelled with mCherry; scale bar, 25 μm). White arrows indicate cells of interest; asterisks indicate foci formation. (**h**) Quantification of G. (*N* > 1000 tracked cells).

## Discussion

Mammalian directed evolution aims to generate biomolecules that are optimized *in situ* for a desired activity. This is dependent on a high mutation rate to diversify the target and a robust link between function and fitness to maintain the integrity of the system, technical challenges that have not yet been overcome. To this end, we have developed PROTEUS, a novel DE platform based on a chimeric SFV design, which takes advantage of the high error rate inherent in alphavirus replication while solving major issues with previous strategies^19^.

Natural viral infections generate a diversity of virions encoding incomplete genomes that collectively compete with host defenses^37^. Our work was motivated by the recent observation that capsid-genome interactions are distributed throughout the alphavirus genome^21^, which contribute to the production of non-functional “cheater” particles containing incomplete genomes that encode structural proteins and contaminate DE campaigns^19^. The capsidless VLVs used in PROTEUS utilize the cellular exosome pathway to escape the host cell^24^, eliminating the co-evolved and highly redundant links between the viral genome and its cognate capsid. This approach maintained system integrity and enabled long-term propagation (>30 rounds) on a synthetic DE circuit. These results pave the way for the adaptation of PROTEUS to other mammalian cell types to provide tissue- or disease-specific environments for DE campaigns.

Our study does have limitations. The mutational bias we observed is consistent with ADAR-mediated editing of SFV RNA genome^38^. It may be necessary to knockout ADAR activity in host cells to neutralize the mutation spectrum. Also, VSVG expression must exceed a certain threshold to support VLV propagation. The circuit activity can be precisely tuned when selective pressure is controlled by a small molecule (e.g. dox), however, it may be more challenging to maintain this balance for more complex targets (e.g. receptors, ligands, signaling molecules) or with more complex circuit designs. Finally, while improved prokaryotic DE systems enable shorter campaigns with very high mutation rates^39^, PROTEUS provides unique utility for targets where a mammalian cellular context is critical, such as chromatin regulators.

The effectiveness of DE campaigns using VLVs depends on how tightly the transgene function is coupled to VLV fitness. We confirmed that PROTEUS generates authentic evolution products by recovering well-characterised doxycycline resistance mutations in tTA. While single mutations appeared in the VLV population at low frequency, they were rapidly outcompeted by double mutant variants that exhibited greater dox resistance, highlighting the power of PROTEUS to generate complex mutations that would be difficult to generate through existing technologies. For example, sampling the complete double mutant sequence space of tTA would require testing 20^247^ combinations, which is beyond any existing experimental approach. In a more challenging test, the sensitivity of the highly optimized rtTA-3G protein was further increased by long-term propagation on limiting amounts of doxycycline. This shows that selective pressure can be maintained over extended DE campaigns. Finally, by optimizing the intracellular function of a nanobody, we generated a p53 biosensor that responds to DNA damage. We anticipate that PROTEUS will be suitable for the generation or optimization of diverse biomolecules designed to function in complex mammalian systems.

## Supporting information

Supplementary Table 1. eGFP-LUC variants.

Supplementary Table 2. tTA variants.

Supplementary Table 3. Analysis of tTA and rtTA-3G mutants.

Supplementary Table 4. rtTA-3G variants.

Supplementary Table 5. Nb139 variants.

Supplementary Table 6. List of plasmids used or generated in this study.

Supplementary Table 7. List of PROTEUS primers.

Supplementary Movie 1. Videos for stills of timelapse in Fig. 4G and Fig S6.

Supplementary Movie 2. Videos for stills of timelapse in Fig. 4G and Fig S6.

## Funding

Tour de Cure Mid-Career Researcher Grant 2023 G219180 (AJC)

The Centenary Institute (DH)

National Health and Medical Research Council GNT2029754 (DH, AJC)

National Health and Medical Research Council GNT1185002 (DH, GGN)

National Health and Medical Research Council GNT1107514 (GGN)

National Health and Medical Research Council GNT1158164 (GGN)

National Health and Medical Research Council GNT1158165 (GGN)

NSW Ministry of Health (DH, AJC, GGN)

Kind donation from Dr. John and Anne Chong (GGN)

## Author contributions

Conceptualization: AJC, CED, DH, GGN

Methodology: AJC, CED, DH, GGN

Investigation: AJC, CED, CLM, TD

Visualization: AJC, CED, DH, GGN

Funding acquisition: DH, GGN

Project administration: DH, GGN

Supervision: DH, GGN

Writing – original draft: AJC, CED, DH, GGN

Writing – review & editing: AJC, CED, DH, GGN

## Data and materials availability

The authors confirm that the data supporting findings of this study are available within the article and its supplementary materials. Basecalled long-read nanopore sequencing FASTQ reads and raw short-read Illumina sequencing FASTQ reads have been deposited at the Gene Expression Omnibus (GEO; GSE250502). PROTEUS plasmids will be made available on Addgene for academic use.

