## Supplemental Materials for "A chimeric viral platform for directed evolution in mammalian cells"

**The PDF file includes:**

Materials and Methods
Extended Data Figs. 1 to 7
Descriptions of Supplementary Tables 1 to 7

**Other Supplementary Materials for this manuscript include the following:**

Supplementary Tables 1 to 7
Supplementary Movies 1 to 2

#### Materials and Methods

##### Cell Culture

BHK-21 [C-13] cells (#CCL-10) were sourced from the American Type Culture Collection (ATCC). Cells were maintained in a humidified 37°C (5% $CO_2$) atmosphere in MEM α (ThermoFisher, #32571101) supplemented with 5% fetal bovine serum (FBS) (Bovogen, #SFBS-F, French origin) and 10% tryptose phosphate broth (TPB) (ThermoFisher, #CM0283B), hereafter defined as 'BHK-21 Growth Medium'. Fresh aliquots of cells were thawed for directed evolution campaigns once in-house passage counts reached 28.

##### Molecular Biology and Plasmid Construction

All PROTEUS system and synthetic circuit plasmids were designed in SnapGene® (Version 6.1). Sequences of interest (e.g. promoter sequences, transgene inserts, plasmid vector backbones etc.) were isolated by high-fidelity PCR amplification with Velocity DNA Polymerase (Bioline, #BIO-21099) or Q5® High-Fidelity 2X Mastermix (NEB, #M0492) using primers synthesized by Integrated DNA Technologies (IDT). Assembly of PCR amplicons was performed using the NEBuilder HiFi DNA Assembly Master Mix (NEB, #E2621) following manufacturer's recommendations. Assembled products were transformed into NEB® 10-beta Competent *E. coli* (NEB, #C3019) and selected on LB agar plates (ThermoFisher, #22700025) supplemented with 100 μg/ml ampicillin (Sigma-Aldrich, #A9518). Single colonies grown overnight in liquid LB broth (ThermoFisher, #12795-084) supplemented with 100 μg/ml ampicillin were processed with the ISOLATE II Plasmid Mini Kit (Bioline, #BIO-52057) for sequence verification. Plasmid constructs were verified by restriction digestion and Sanger sequencing at the Australian Genome Research Facility (AGRF). Once successful assembly had been confirmed, fresh streaks of confirmed positive constructs were generated on antibiotic-supplemented LB agar plates and cultures generated for plasmid isolation using the PureYield™ Plasmid Maxiprep System (Promega, #A2393), producing the yields and purity necessary for transfection applications. Plasmids used or generated in this study are listed in **Supplementary Table 6**.

*PROTEUS Virus-Like Vesicle (VLV) SFV Constructs*

The initial SFV DNA plasmid construct (pSFV-DE_eGFP-LUC) was designed for expression from a constitutive CMV promoter in the mammalian expression vector pcDNA3.1(+). This parental plasmid was linearized by PCR and the amplicon purified by gel extraction. SFV elements (5'UTR, NSP1-4, 3'UTR), the sequences of which were derived from pSFV3 (Addgene #92072) (*28*), were synthesized as a series of gBlocks HiFi Gene Fragments by IDT and used for NEBuilder HiFi assembly. We built fourteen point mutations into the NSP genome sequence which evolved in a capsid-deficient SFV strain for production of high-titer VLVs enveloped with VSVG (*15*). Two intermediates containing separate halves of the NSP1-4_eGFP-LUC_3'UTR insert were generated from which the final assembly was obtained by PCR and HiFi assembly.

Subsequently, the NSP2 $^{674}$ADA$^{676}$ codon Vloop was mutated to $^{674}$RLE$^{676}$ by HiFi assembly to attenuate cytopathic effects (*29*). This attenuated VLV (pSFV-DE) forms the basis of the SFV DNA replicon system presented within this study, and all transgenes for directed evolution were inserted as a direct replacement of the eGFP-LUC coding sequence by PCR and HiFi assembly.

**Packaging and Amplification of PROTEUS VLVs**

*Packaging (RP)*

PROTEUS VLVs were packaged in BHK-21 cells following transfection with plasmid DNA. BHK-21 cells were seeded in 6-well plates at a density of 1.95 x $10^5$ cells/well in 2 mL BHK-21 Growth Medium and incubated for 24 hours. A total of 1 μg of plasmid DNA (2:1:1 of pCMV_VSVG, pSFV-DE_[transgene] and pSFV-DE_eGFP-LUC) was diluted in Opti-MEM Reduced Serum Medium with GlutaMAX Supplement (ThermoFisher, #51985034) and transfected into cells using *Trans*IT-2020 Transfection Reagent (Mirus Bio, #MIR5400) following the manufacturer's recommendations (employing a 3:1 transfection reagent:DNA ratio). 6 hours post-transfection, cells were rinsed twice with DPBS (Sigma-Aldrich, #D8537) and 1.2 mL of BHK-21 Growth Medium was added. VLV-containing supernatant was collected 24 hours post-transfection and centrifuged at 1000 g for 5 minutes to pellet cellular debris. Clarified supernatants were collected for titration and subsequent transduction experiments.

*Amplification (RA)*

Packaged VLVs from RP were propagated further in constitutive VSVG-expressing BHK-21 cells to amplify infectious, enveloped particles. BHK-21 cells were seeded in 6-well plates (1.95 x $10^5$ cells/well in 2 mL) in BHK-21 Growth Medium and incubated for 24 hours. A total of 1 μg of pCMV_VSVG DNA was transfected into cells using *Trans*IT-2020 Transfection Reagent. 6 hours post-transfection, cells were rinsed twice with DPBS and neat RP VLVs added (500 μL for 6-well plates) in the presence of 8 μg/ml polybrene (Sigma-Aldrich, #H9268). Cells were incubated with VLVs for 1 hour and rinsed twice with DPBS before 1.2 mL BHK-21 Growth Medium was added for a further 23 hours of incubation. VLV-containing supernatants were collected and processed as described in *Packaging (RP)*.

**VLV Titration**

VLV-containing supernatants were titrated as per (*11*) using an NSP2-specific primer-probe set. Following collection and clarification, undiluted VLV-containing supernatants were combined with the TaqMan™ Fast Virus 1-Step Master Mix (ThermoFisher, #4444434) in a minimum of technical duplicates. Plates were run on a QuantStudio™ 7 Flex or QuantStudio™ 6 Pro Real-Time PCR System (ThermoFisher) using 'Fast' protocol parameters. Serially diluted pSFV_eGFP-LUC plasmid DNA was used to generate a standard curve for absolute quantification (ranging between $10^3$-$10^7$ genome copies (gc) per reaction). Plotted standard curves were used to determine VLV titers in gc/mL. Detection thresholds were determined with no template control reactions and typically ranged between $10^2$-$10^3$ gc/mL. For subsequent VLV transductions, these values were subtracted from calculated titers. Primer sequences are listed in **Table S7**.

**VLV Transduction (Evolution Round 1 (E1)-onwards)**

For evolution, amplified VLVs from RA were propagated in synthetic circuit-expressing cells that require VLV-encoded transgene functionality to activate transcription of the VSVG packaging element. BHK-21 cells were seeded in 6-well plates at a density of 1.95 x $10^5$ cells/well in 2 mL BHK-21 Growth Medium, incubated for 24 hours, and transfected using *Trans*IT-2020 Transfection Reagent (Mirus Bio, #MIR5400) following the manufacturer's recommendations with a total of 1 μg of synthetic circuit-encoding plasmids. The SRF circuit used 1 μg of pSRE_VSVG; the tTA/rtTA circuits used 1 μg of pTRE3G_VSVG; the Nb139 circuit supplied two plasmids at a 1:1 ratio (p53-ZF43:pZF43-VSVG), 0.5 μg/plasmid. Growth medium was replaced prior to transfection if the circuit of interest required chemical additives or adjusted media (e.g. reduced [FBS]). 6 hours post-transfection, a mock-transfected well was trypsinized and counted to calculate the volume of titered VLV inoculum needed to achieve an MOI of 1 gc/cell. Typical counts ranged between 3-10 x $10^5$ cells/well. An MOI of 1 is used to restrict circuit activation to the activity of a single transgene variant per cell to minimize crosstalk. Cells were rinsed once with DPBS before VLV was applied in a 500 µL volume of BHK-21 Growth Medium supplemented with 8 µg/ml polybrene. Cells were incubated with VLVs for 1 hour and rinsed twice with DPBS before 1.2 mL BHK-21 Growth Medium was added for a further 23 hours of incubation (supplemented with chemical additives (e.g. doxycycline or reduced [FBS] where relevant)). VLV-containing supernatants were collected and processed as described in *Packaging (RP)*.

Each subsequent round uses the titered VLVs produced in the preceding round to iteratively diversify, select and amplify variants of improved fitness.

Notes: the number of cells transduced, and hence the number of variant transgene-carrying VLVs screened per round, can be increased by proportionally upscaling to larger flask footprints. All evolution campaigns presented within this article represent experiments performed in 6-well plates. An MOI >1 could be used to achieve higher circuit activation, but dominance of high fitness mutations may be delayed by the piggybacking of low fitness variants following co-transduction of a single cell. The concentration of dox used to initiate tTA and rtTA campaigns was optimized to permit propagation of VLVs.

**Luciferase/Resazurin Assay**

BHK-21 cells were seeded in 2 x 96-well plates at 6.6 x $10^3$ cells/well. Cells were transfected with a total 34 ng of plasmid constructs using *Trans*IT-2020 Transfection Reagent following the manufacturer's recommendations. At 6 hours post-transfection, cells were washed twice with DPBS, fresh BHK-21 Growth Medium added and the plates incubated for 24 hours. Cell viability (as a proxy for cell density) was determined by replacement of growth medium with 30 µg/ml resazurin (Sigma-Aldrich, #R7017)-supplemented medium and incubation for approximately 30 min. Resorufin fluorescence was measured on an Infinite M1000 PRO microplate reader (Tecan). Supplemented medium was removed before cells were rinsed once with room temperature DPBS and replaced with 30 µL growth medium. Luciferase activity was assessed using the Steady-Glo® Luciferase Assay System (Promega, E2510) following manufacturer's recommendations in black-bottom plates. Firefly luminescence was measured on an Infinite M1000 PRO microplate reader (Tecan). Raw firefly luminescence values were background-subtracted and then normalized against well-matched resazurin fluorescence prior to analyses.

**Transgene Isolation**

SFV RNA was extracted from 140 µL culture supernatant using the QIAamp Viral RNA Mini Kit (Qiagen, #52904) following the manufacturer's recommendations. SFV RNA was reverse transcribed into cDNA and all SFV transgene sequences, positioned between NSP4 and the viral 3' untranslated region (UTR), were PCR-amplified using the OneTaq® One-Step RT-PCR Kit (E5315S). PCR was conducted using an NSP4-specific forward primer and a viral 3'UTR-specific reverse primer (listed in **Supplementary Table 7**). Amplified transgenes were resolved by gel electrophoresis using standard protocols and amplicons of the appropriate size range for the transgene of interest were purified by gel extraction using the ISOLATE II PCR and Gel Kit (Bioline, #BIO-52060). Amplicons were sequenced through long-read nanopore, short-read

Illumina, or single read Sanger sequencing approaches. For Sanger sequencing, amplicon transgene DNA was blunt cloned into the linear plasmid backbone of the PCR Cloning Kit (NEB, #E1202S) using manufacturer's recommendations. Following transformation, single colonies were cultured and sequenced as described in **Molecular Biology and Plasmid Construction**.

**Long-read (Oxford Nanopore) Sequencing**

*Sample Processing*

RT-PCR-isolated transgene DNA sequences were processed to generate the libraries required for full-length transgene sequencing with Oxford Nanopore Technologies (ONT) Flongle flow cells (ONT, #FLO-FLG001). Sequencing samples were prepared following the recommendations found within the ONT protocol 'Amplicons by Ligation' (version ACDE_9110_v110_revV_10Nov2020). DNA concentrations were determined using the Qubit™ dsDNA HS Assay Kit (ThermoFisher, #Q32851) and 200 fmol of DNA was processed using the NEBNext Companion Module for ONT Ligation Sequencing (NEB, #E7180). Sequencing adapters were ligated onto sequences with the Ligation Sequencing Kit (ONT, #SQK-LSK110).

*Sequencing and Basecalling*

Up to 40 fmol of each DNA library was loaded on ONT Flongle flow cells (R9.4.1 chemistry, #FLO-FLG001) in a MinION Mk1B Sequencer (ONT, #MIN-101B) fitted with a Flongle Adapter (ONT, #ADP-FLG001). Sequencing was performed using MinKNOW (ONT, version 4.2.8) using default parameters and the following inputs: kit used, SQK-LSK110; 0.5 hours between MUX scans; basecalling, disabled. A minimum $2 \times 10^4$ raw reads were obtained per flow cell. Raw FAST5 files were basecalled using Guppy (version 4.5.2) with the minimum q-score filter set to 7.0. Basecalled FASTQ reads have been deposited at the Gene Expression Omnibus (GEO; GSE250502).

*Alignment*

Quality-filtered basecalled reads (in FASTQ format) were processed using EPI2ME Desktop Agent (ONT, version 3.3.0.1031). Sequence reference files were uploaded into the program with the Fasta Reference Upload workflow (v2021.07.15). For each sample, all reads were aligned to reference files using the Fastq Custom Alignment workflow (v2021.03.25) using default parameters.

**Short-read (Illumina) Sequencing**

*Sample Processing*

RT-PCR-isolated transgene DNA sequences were processed further to generate the yield of DNA required for short-read Illumina sequencing with NovogeneAIT Genomics Singapore (Novogene). Extracted DNA was further amplified by high-fidelity PCR amplification with Q5® High-Fidelity 2X Mastermix (NEB, #M0492) using the same primer set as for initial transgene isolation. Amplicons were resolved by DNA gel electrophoresis and purified to have a minimum 1.5 µg for sequencing.

*Sequencing*

Samples were sequenced with Novogene using their 'Microbial PCR Product Whole Genome Sequencing' strategy. Amplicons underwent fragmentation prior to PCR-free library preparation and sequencing using NovaSeq PE250 technology. Raw FASTQ reads have been deposited at the Gene Expression Omnibus (GEO; GSE250502).

*Variant Nucleotide Analysis*

To identify variant nucleotides, raw FASTQ files were aligned to transgene reference sequences (FASTA) with *minimap2*. Any mate-pair issues were fixed in the output .sam files with the *samtools fixmate* command before reads were sorted (*samtools sort*) and reformatted (*samtools mpileup*). VarScan 2.3.9 (*30*) was used to call for variants with command *mpileup2snp* with arguments *--min-coverage 10000 --min-reads2 15 --min-var-freq 0.00025 --p-value 0.01*. Further annotation was performed with *vcf-annotator* using default commands, aligning to transgene reference sequences (GenBank format).

**Immunocytochemistry**

Cells were fixed with 4% PFA for 20 minutes at room temperature. Blocking was performed in PBS with 5% Normal Goat Serum, 1% BSA, 0.05% Triton-X, 0.3M Glycine for 1 hour at room temperature. Cells were then incubated with 1:400 anti-P53 mAb (Clone 7F5) (Cell Signaling Technology, #2527) in blocking solution for one hour at room temperature, followed by five washes with PBS-0.05% Triton-X. A goat anti-rabbit Alexa647 secondary antibody (Invitrogen, #A21245) and Hoechst 33142 in blocking solution were then used to incubate cells for one hour at room temperature. Cells were then washed 5 times with PBS-0.05% Triton-X, and imaged in PBS.

**Epifluorescence Microscopy**

Phase contrast and eGFP fluorescence images were obtained on an Axio Vert.A1 FL (Zeiss) microscope fitted with an AxioCam ICM1 camera (Zeiss 60N-C 2/3" 0.63X adapter) at 5X magnification. eGFP images were captured with a BP475/40 excitation and BP530/50 emission filter (FT500 beam splitter). Images were collected with Zen 2 Blue Edition (Zeiss, version 2.0.0.0).

**High-throughput Imaging**

*Fixed*

Cells were imaged using the Opera Phenix Plus High-Content Screening system. High-throughput image analysis was performed using Harmony Software V5.1 (Perkin Elmer).

*Live*

Cells were imaged using the Opera Phenix Plus High-Content Screening system. Live cell tracking was performed by detecting nuclear fluorescence across a period of 24 hours (~11 min intervals) using Harmony Software V5.1 (Perkin Elmer). Tracked objects with detectable nuclear GFP signal were exported.

**Protein Structure Prediction**

The AlphaFold2_mmseqs2 Google Colab notebook from ColabFold (v1.5.2-patch; https://colab.research.google.com/github/sokrypton/ColabFold/blob/main/AlphaFold2.ipynb; https://github.com/sokrypton/ColabFold) was used to predict protein structures using default settings. The top-ranked prediction by average pLDDT was used for annotation and visualization with UCSF Chimera (Version 1.17.3), developed by the Resource for Biocomputing,

Visualization, and Informatics at the University of California, San Francisco, with support from NIH P41-GM103311 (*31*).

**Statistics**

Statistical analyses were performed in GraphPad Prism 9.2.0, GraphPad Software, San Diego, California USA, www.graphpad.com. All data were plotted as mean ± SEM. Cisplatin treatment assays, VLV titer plots and AF plots were analyzed with two-tailed unpaired t-tests. Fold-changes in luciferase activity were statistically analyzed with a non-parametric Kruskal-Wallis test with a Dunn's multiple comparisons test (**Fig. 4b**) or a repeated measures one-way ANOVA with the Geisser-Greenhouse correction with a Dunn's multiple comparisons test (**Fig. 4f**). Means were compared to the control baseline mean. Changes in nuclear eGFP over time were analyzed using a two-way ANOVA with Šídák's multiple comparisons test. *p*-values<0.05 were considered significant (*, $p<0.05$; **, $p<0.01$; ***, $p<0.001$; ****, $p<0.0001$).

#### Extended Data Fig. 1

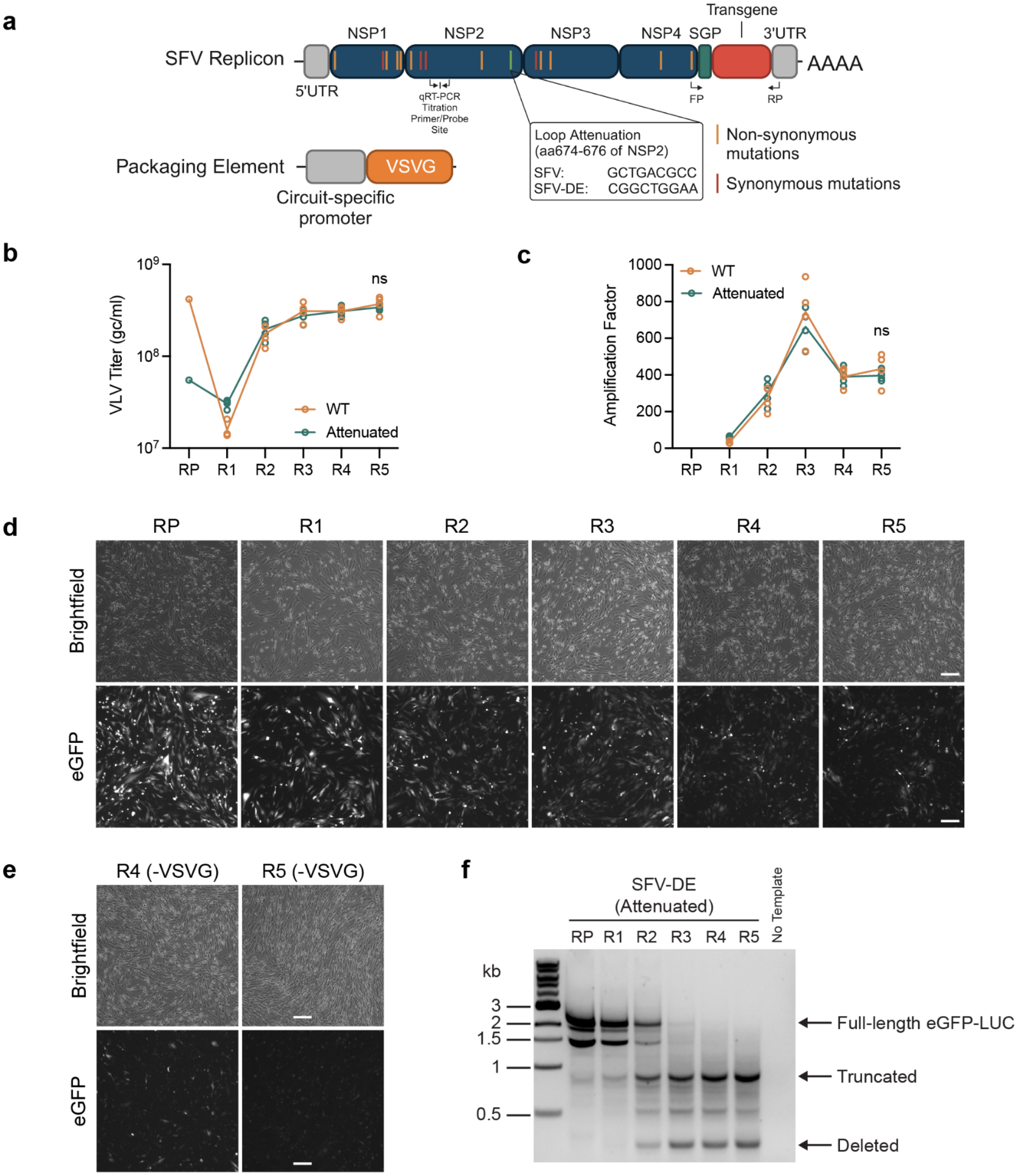


**Extended Data Fig. 1. Benchmarking the PROTEUS platform. (a)** Schematic representation of the PROTEUS SFV DNA replicon genome. FP/RP indicate the positions of the primers used for transgene isolation/sequencing. Propagation **(b)** and amplification **(c)** of WT and ADA>RLE attenuated eGFP-LUC VLVs in CMV_VSVG-expressing BHK-21 cells (*N* = 4). **(d)** Brightfield and epifluorescence microscopy of +VSVG BHK-21 cells at RP or following transduction with eGFP-LUC VLVs at RA-R4 (representative of *N* = 4). Scale bars, 200 µm. **(e)** Microscopy of -

VSVG BHK-21 cells transduced at R3 and R4 (representative of *N* = 4). Scale bars, 200 µm. **(f)** RT-PCR and DNA gel electrophoresis of SFV-DE eGFP-LUC VLV transgenes (*N* = 4 pooled).

### Extended Data Fig. 2

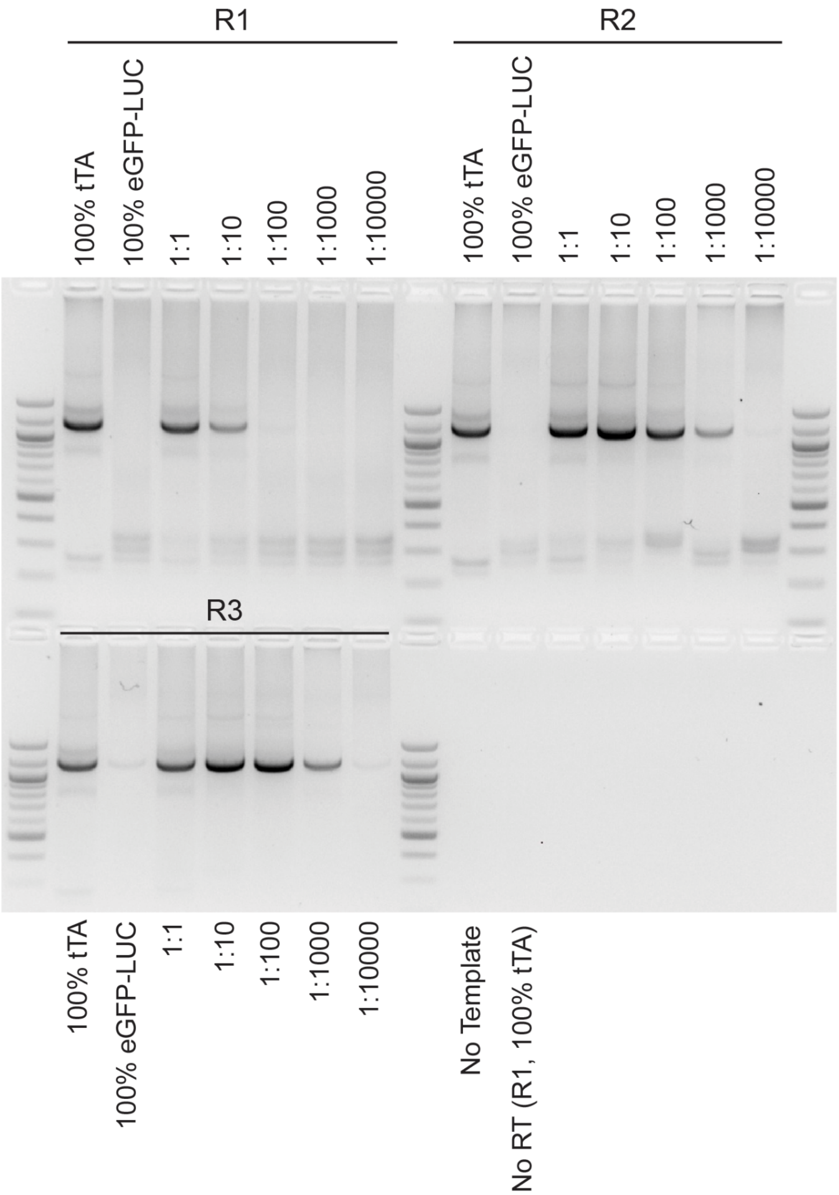


**Extended Data Fig. 2. RT-PCR and DNA gel electrophoresis of isolated transgenes from serially diluted SFV-DE tTA : eGFP-LUC VLVs (R1-R3; *N* = 3 pooled).**

### Extended Data Fig. 3

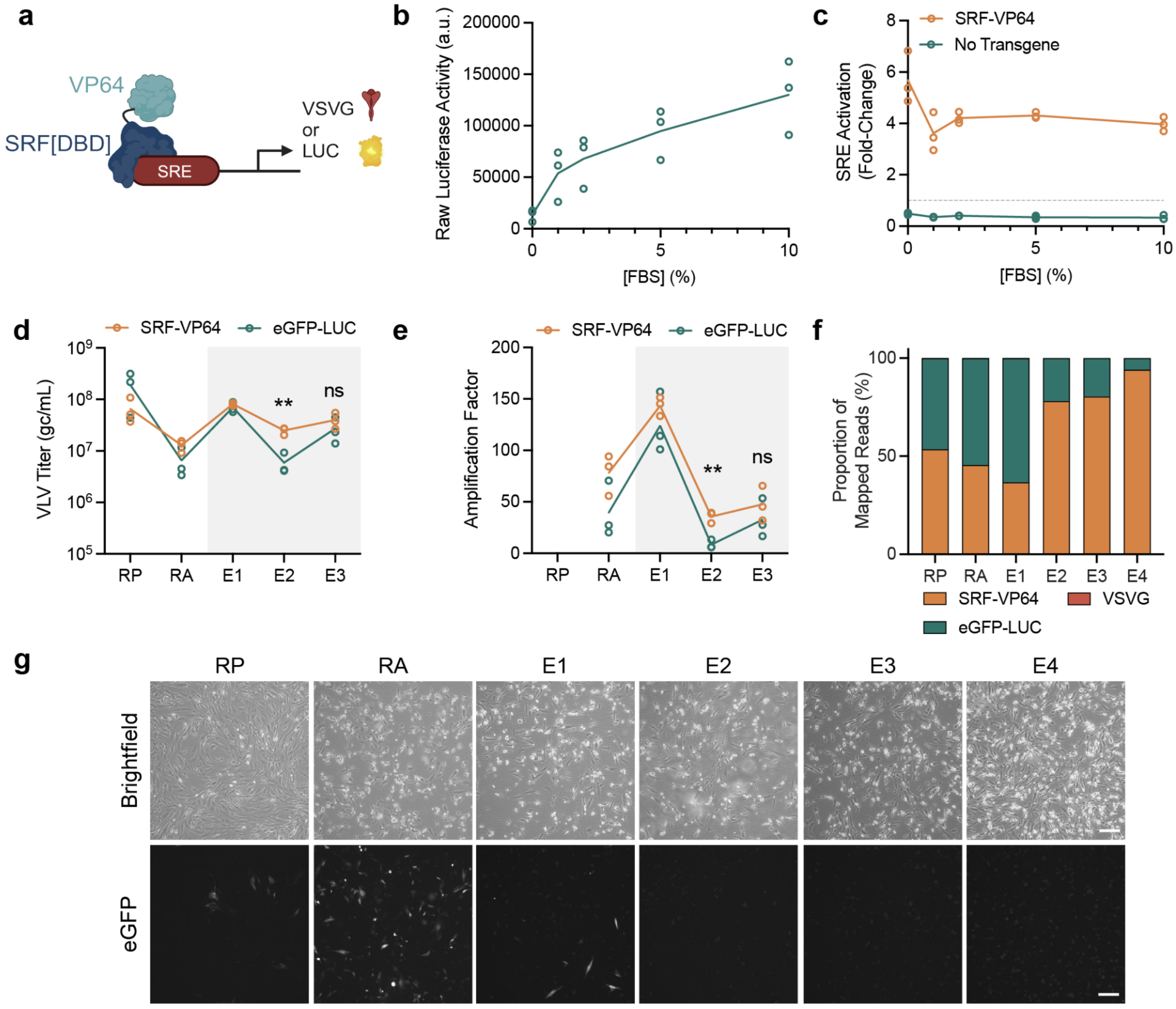

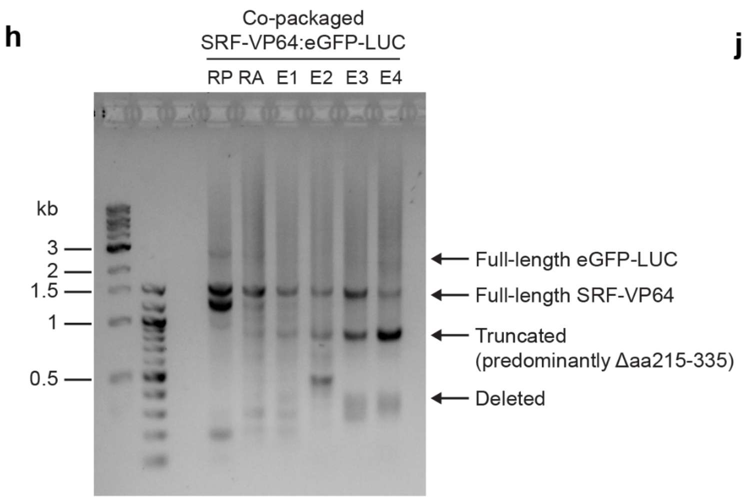


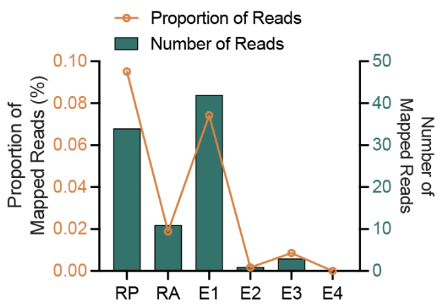


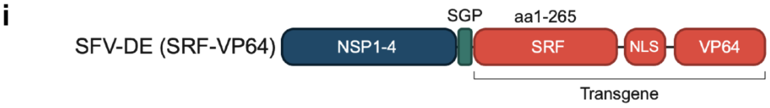


| Protein Domain | Amino Acid Substitution | Nucleotide Substitution | Frequency (%) E3 | Frequency (%) E4 |
|---|---|---|---|---|
| SRF | RGS(aa14-16)> SGQRL and Δaa17-136 | AGAGGCT>**TCTGGGC**AG AGGCTG (bold is inserted) AND deletion of coding sequence for Δaa17-136 | 81.82 | 90.91 |
| | A6A (synonymous) | T>A | 9.09 | N.D. |
| | Q182R | A>G | N.D. | 18.18 |
| | A198A (synonymous) | C>T | N.D. | 9.09 |
| | R200R (synonymous) | G>A | N.D. | 9.09 |
| | C216R | T>C | 9.09 | N.D. |
| | N220D | A>G | N.D. | 9.09 |
| VP64 | D5D (synonymous) | T>C | 9.09 | N.D. |
| | M10T | T>C | N.D. | 9.09 |
| | Sequence identified as derived from eGFP | N/A | 9.09 | 9.09 |

**Extended Data Fig. 3. VLV propagation is dependent on host expression of VSVG. (a)** Circuit design for SRF-VP64 mediated activation of SRE. **(b)** FBS dose-dependent SRE_LUC reporter activation by endogenous factors ($N = 3$). **(c)** SRF-VP64-mediated induction of an SRE-regulated LUC reporter ($N = 3$). Titers **(d)** and amplification factors **(e)** of SRF-VP64 and neutral eGFP-LUC VLPs propagated on cells expressing VSVG under the control of an SRE promoter in 1% FBS-supplemented growth medium (gray box) ($N = 3$). **(f)** Nanopore sequencing of transgene RNA isolated from pooled co-packaged VLP samples ($N = 6$) aligned to reference sequences (>10,000 reads per sample). For E1-E4, VLPs were propagated in 1% FBS-supplemented growth medium. **(g)** Brightfield and epifluorescence microscopy of BHK-21 cells at RP or following transduction with a 1:1 packaged cohort of SRF-VP64:eGFP-LUC VLVs at RA-E4 (representative of $N = 6$). Scale bars, 200 μm. **(h)** RT-PCR and DNA gel electrophoresis of isolated transgenes from a 1:1 packaged cohort of SRF-VP64:eGFP-LUC VLVs ($N = 6$ pooled). (**i**) The DNA bands from E3 and E4 marked as 'Truncated' in (**h**) were cloned and sequenced (n=11 clones per sample). **(j)** Reads from (**f**) (>10,000 per sample) were aligned to a VSVG reference sequence. For E1-E4, VLVs were propagated in 1% FBS-supplemented growth medium.

### Extended Data Fig. 4

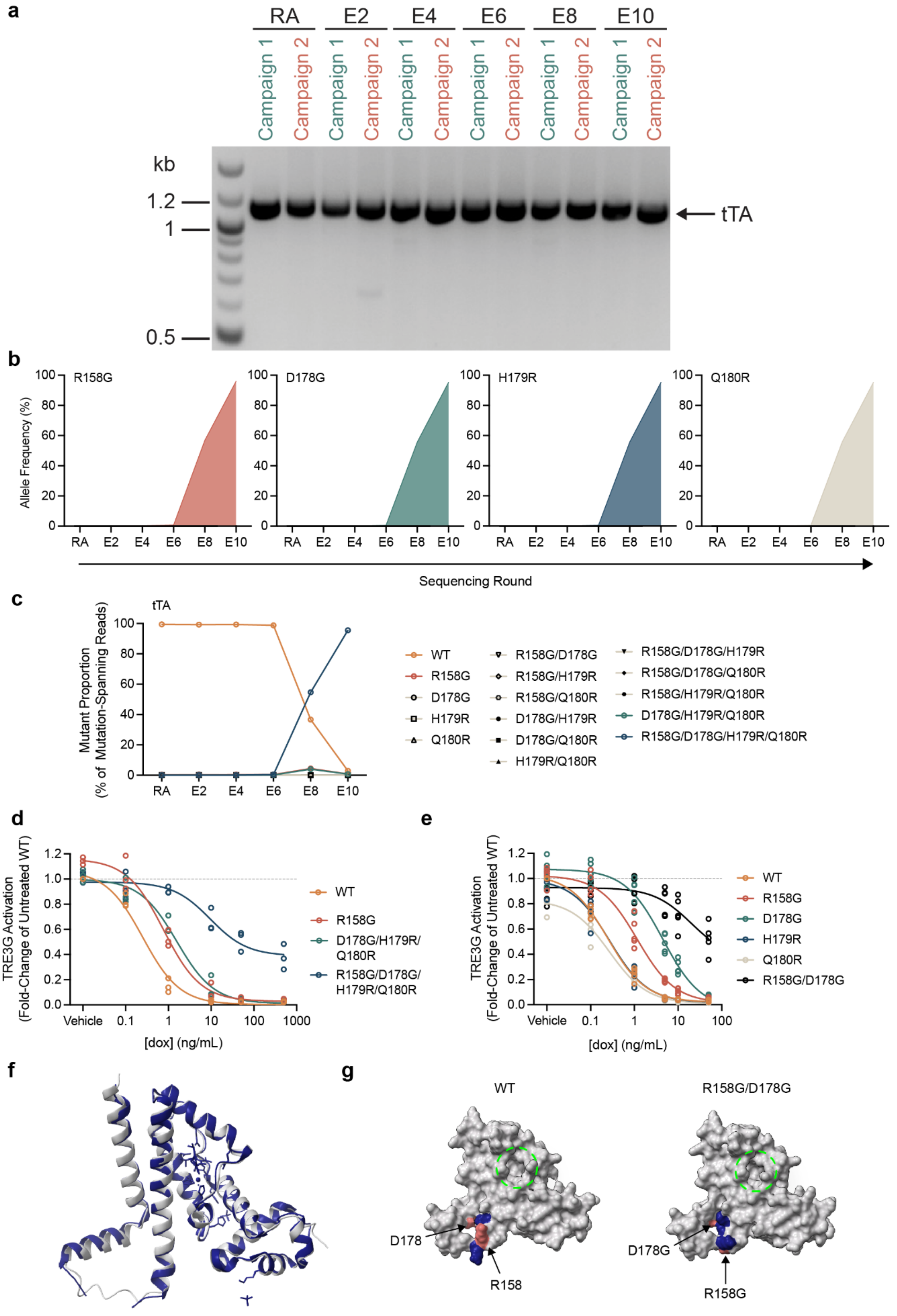

**Extended Data Fig. 4. Directed evolution of tTA (Campaign 2). (a)** DNA gel electrophoresis of isolated tTA transgenes following RT-PCR (from independent Campaigns 1 and 2). Allele frequency of the major variants individually (**b**) and combined (**c**) identified in Campaign 2. A similar analysis of mutation combinations for Campaign 1 cannot be extracted from short-read Illumina sequencing because of the distance between the identified Q32R and R158G residues. Note, a synonymous passenger mutation (E159E) rose to similar levels as each of the R158G, D178G, H179R, Q180R mutations during this campaign. Analysis in (c) permitted synonymous substitutions at E159 to capture all reads analyzed in (b). (**d**) Dox-resistance of evolved tTA variants ($N = 5$). (**e**) Isolated effects of aa178-180 single mutants ($N = 5$). (**f**) Alignment of crystal structure 4AC0 (blue) with Alphafold2-modeled tTA (gray). (**g**) Variant-induced structural changes in tTA modeled with AlphaFold2 (red, mutated residues; blue, displaced functional groups; dashed green circle, drug binding pocket).

### Extended Data Fig. 5

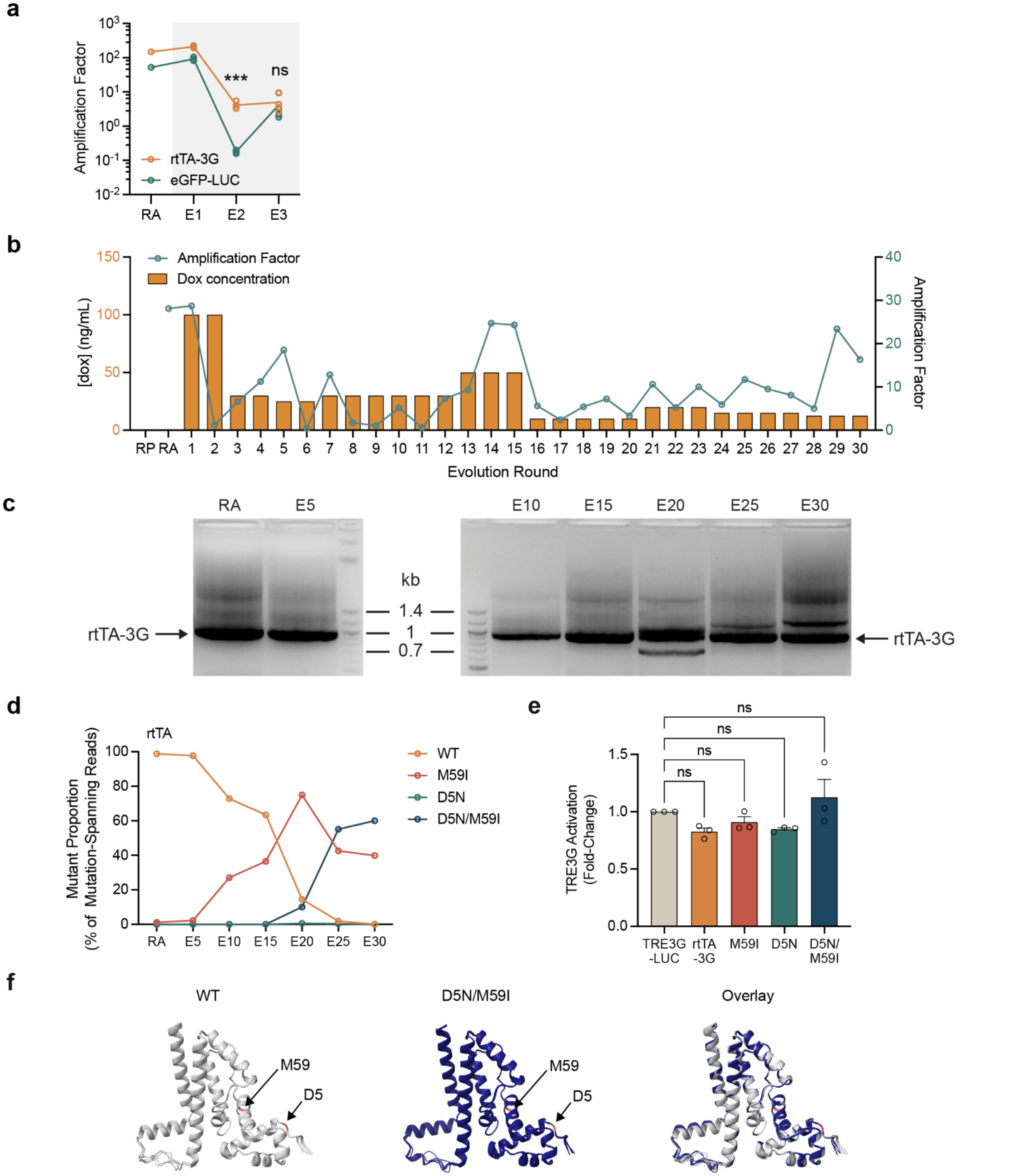


**Extended Data Fig. 5. Directed evolution of rtTA-3G.** (**a**) Amplification factors of VLVs propagated on cells expressing VSVG under the control of TRE3G at 100 ng/mL dox (gray box) ($N$ = 4). (**b**) Dox concentration and AF for each round of rtTA-3G evolution. (**c**) DNA gel electrophoresis of isolated rtTA transgenes following RT-PCR. (**d**) Allele frequencies of single and double mutant variants during long-term propagation on minimal concentrations of dox. (**e**)

Basal activity of the evolved variants in the absence of dox ($N = 3$). (**f**) Alignment of top 5 ranked Alphafold2 predictions for rtTA-3G (tan) and the D5N/M59I variant (blue; red, mutated residues).

### Extended Data Fig. 6

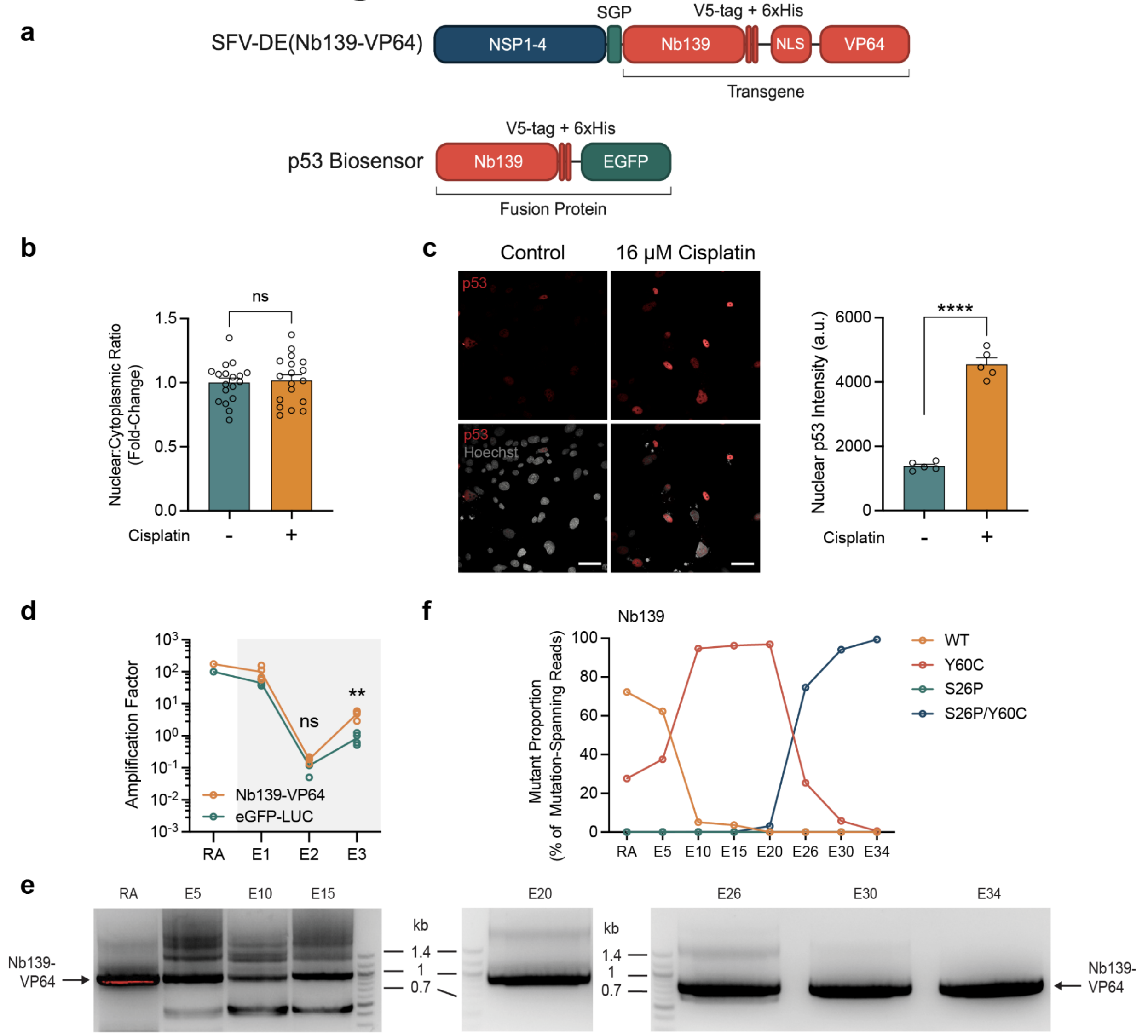


**Extended Data Fig. 6. Directed evolution of a nanobody with PROTEUS.** (**a**) Schematic of Nb139-VP64 and Nb139-eGFP fusions. (**b**) Effect of 16 µM cisplatin on parental Nb139-eGFP nuclear localization ($N$ = 18). (**c**) p53 immunofluorescence (red) and Hoechst nuclear staining (gray); Quantification of nuclear p53 accumulation. Scale bars, 50 µm. ($N$ ≥ 272 cells analyzed per replicate). (**d**) Amplification factors of VLVs propagated on cells expressing VSVG under the control of a p53 2-hybrid circuit (gray box) ($N$ = 4). (**e**) DNA gel electrophoresis of isolated Nb139-VP64 transgenes following RT-PCR. (**f**) Allele frequencies of single and double mutant variants during long-term propagation on the 2-hybrid circuit.

### Extended Data Fig. 7

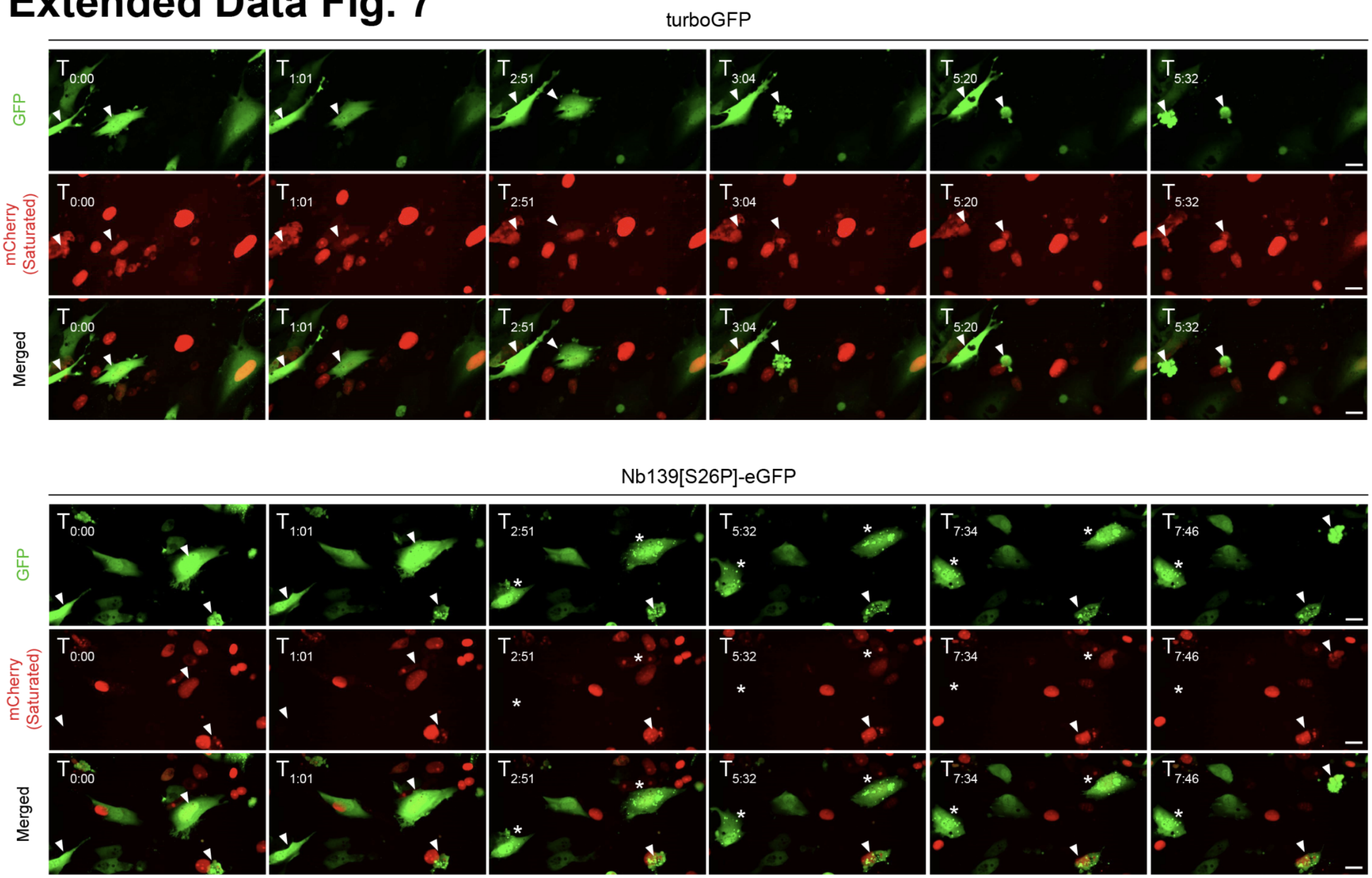


**Extended Data Fig. 7. Individual channels from timelapse in Fig. 4G.** Nuclei labeled with mCherry (red). Nb139[S26P]-eGFP biosensor or turboGFP control (green). White arrows indicate cells of interest; asterisks indicate foci formation. Scale bars, 25 µm.

**Supplementary Table 1. eGFP-LUC variants.** All substitutions in transgene RNA detected at ≥0.3% over three rounds of VLV propagation.

**Supplementary Table 2. tTA variants.** Non-synonymous substitutions in transgene RNA detected at ≥1% during two PROTEUS campaigns.

**Supplementary Table 3. Analysis of tTA and rtTA-3G mutants**. Half maximal inhibitory/effective concentrations (IC50/EC50) and 95% confidence intervals (95% CI) for tTA and rtTA-3G figures.

**Supplementary Table 4. rtTA-3G variants.** Non-synonymous substitutions in transgene RNA detected at ≥1% of the population during a PROTEUS campaign.

**Supplementary Table 5. Nb139 variants.** Non-synonymous substitutions in transgene RNA detected at ≥1% during a PROTEUS campaign.

**Supplementary Table 6. List of plasmids used or generated in this study.**

**Supplementary Table 7. List of PROTEUS primers.**

**Supplementary Movie 1. Videos for stills of timelapse in Fig. 4G and Fig S6.** turboGFP control in green. Nuclei labeled with mCherry (red).

**Supplementary Movie 2. Videos for stills of timelapse in Fig. 4G and Fig S6.** Nb139[S26P]-eGFP biosensor in green. Nuclei labeled with mCherry (red).
